# Hormone Levels Are Related to Altered Functional Connectivity in Prolactinomas

**DOI:** 10.1101/746297

**Authors:** Shun Yao, Chenglong Cao, Pan Lin, Parikshit Juvekar, Ru-Yuan Zhang, Matthew Vera, Ailiang Zeng, Alexandra J. Golby, Guozheng Xu, Yanmei Tie, Jian Song

## Abstract

**Background and Objective:** Prolactinomas may cause drastic hormone fluctuations throughout the body. It is not fully understood how endogenous hormone disorders such as prolactinomas reshape the patient’s brain. By employing the resting-state functional magnetic resonance imaging technique, we aimed to investigate the whole-brain functional connectivity (FC) and its relationship with hormone levels in patients with prolactinomas.

**Methods:** Using whole-brain and seed-based functional connectivity analyses, we compared FC metrics between 33 prolactinoma patients and 31 healthy controls matched with age, sex, and handedness. Then we performed partial correlation analysis to examine the relationship between FC metrics and hormone levels.

**Results:** Compared to healthy controls, we found that prolactinoma patients showed significantly increased thalamocortical (visual cortex) and cerebellar-cerebral connectivity. In addition, endogenous hormone levels were positively correlated with the increased FC, and the hormone-FC relationships showed sex difference in prolactinoma patients.

**Conclusions:** Our findings are the first to reveal the altered FC patterns and sex-dependent hormone-FC relationships in prolactinoma patients, indicating the important role of hormone levels in the neural mechanism of brain reorganization and hyperactive intrinsic connections in prolactinomas.

## INTRODUCTION

Prolactinomas are characterized by a dramatic surge of prolactin that suppresses the secretion of sex steroid hormones. Evidence has been found that sex steroids can affect the structural and functional organization of the brain. We have previously investigated the influence of endogenous hormones on brain gray matter and neurocognition in prolactinoma patients.^1^ However, functional connections in prolactinoma patients have scarcely been explored. Recently, the resting-state functional magnetic imaging (rsfMRI) has become a useful tool to characterize the brain’s functional connectivity (FC) pattern that is considered a potential metric reflecting the temporal dependence of neuronal activation patterns of spatially separated brain areas.^2^

The aim of the current study was to determine the functional connectivity patterns in prolactinomas. We hypothesized that prolactinoma patients would show abnormal functional connectivity patterns related to the increased endogenous hormone levels.

## METHODS

### Study Population

Thirty-three prolactinoma patients and 31 healthy controls participated in this study. All patients were recruited in the Department of Neurosurgery during diagnostic hospitalization. The inclusion and exclusion criteria were described in our previous work^1^.

All procedures were under the Declaration of Helsinki and approved by the Ethical Committee of General Hospital of Chinese PLA Central Theater Command (Approved ID: [2017] 024-1). The study protocol was fully explained, and written informed consent was acquired from all participants.

### Hormone Assays and Visual Assessment

The measurement of hormone levels in serum was described in our previous work^1^. All patients routinely underwent complete ophthalmologic examinations at admission. The E chart was used to measure the best-corrected visual acuity. The visual field parameter was obtained using the standardized, automated perimetry (Octopus 900 Perimetry, Switzerland).

### MRI Data Acquisition

The MRI data were acquired with a 1.5 Tesla scanner (GE EXCITE, Milwaukee, USA) using an 8-channel head coil. The structural MRI acquisition was described in our previous study^1^. Blood oxygen level dependent (BOLD) fMRI data were acquired using echo-planar images (TR/TE 2000/30 ms, 4.0 mm thickness, 0.5 mm gap, 33 axial slices, matrix size of 64 × 64, and 7 min). During rsfMRI, all participants were instructed to keep their eyes open with a central fixation condition.

### fMRI Preprocessing

fMRI data were preprocessed using the CONN Toolbox (v.18.a; http://www.nitrc.org/projects/conn) based on SPM12 (http://www.fil.ion.ucl.ac.uk/spm/software/spm12) in MATLAB R2018a (MathWorks, Inc., MA, USA). Standard preprocessing steps of rsfMRI included realignment, slice-timing correction, functional outlier detection, normalization, Gaussian spatial smoothing (6 mm FWHM), bandpass filtering (0.008-0.09 Hz), and cleaning of motion and physiological noise using the CompCor method.

### Whole-brain Functional Connectivity Analysis

For the whole-brain FC analyses, 132 regions of interest (ROIs) were defined using the brain parcellation implemented in the CONN toolbox. The mean time series of each ROI was extracted to construct the ROI-to-ROI correlation matrices, which were then converted to z scores using Fisher’s z-transformation. To correct multiple comparisons between groups, a seed-level false discovery rate (FDR) method with the significant threshold of *p* < 0.05 was performed.

### Statistical Analysis

Group differences were compared using two-sample student’s t-test or Pearson’s Chi-Squared test, as appropriate. The partial correlation analysis was performed to determine the relationship between FC metrics and hormone levels controlling the factors of age, tumor volume, and disease history. Significance was set at p < 0.05 (two-tailed). Statistical analyses were performed using JASP version 0.10.2 (https://jasp-stats.org/).

## RESULTS

### Clinical Characteristics

There were no differences between prolactinoma patients and healthy control in age (*p* = 0.266) and sex (*p* = 0.477). The other clinical characteristics were described in **Table 1**.

**Table 1.**
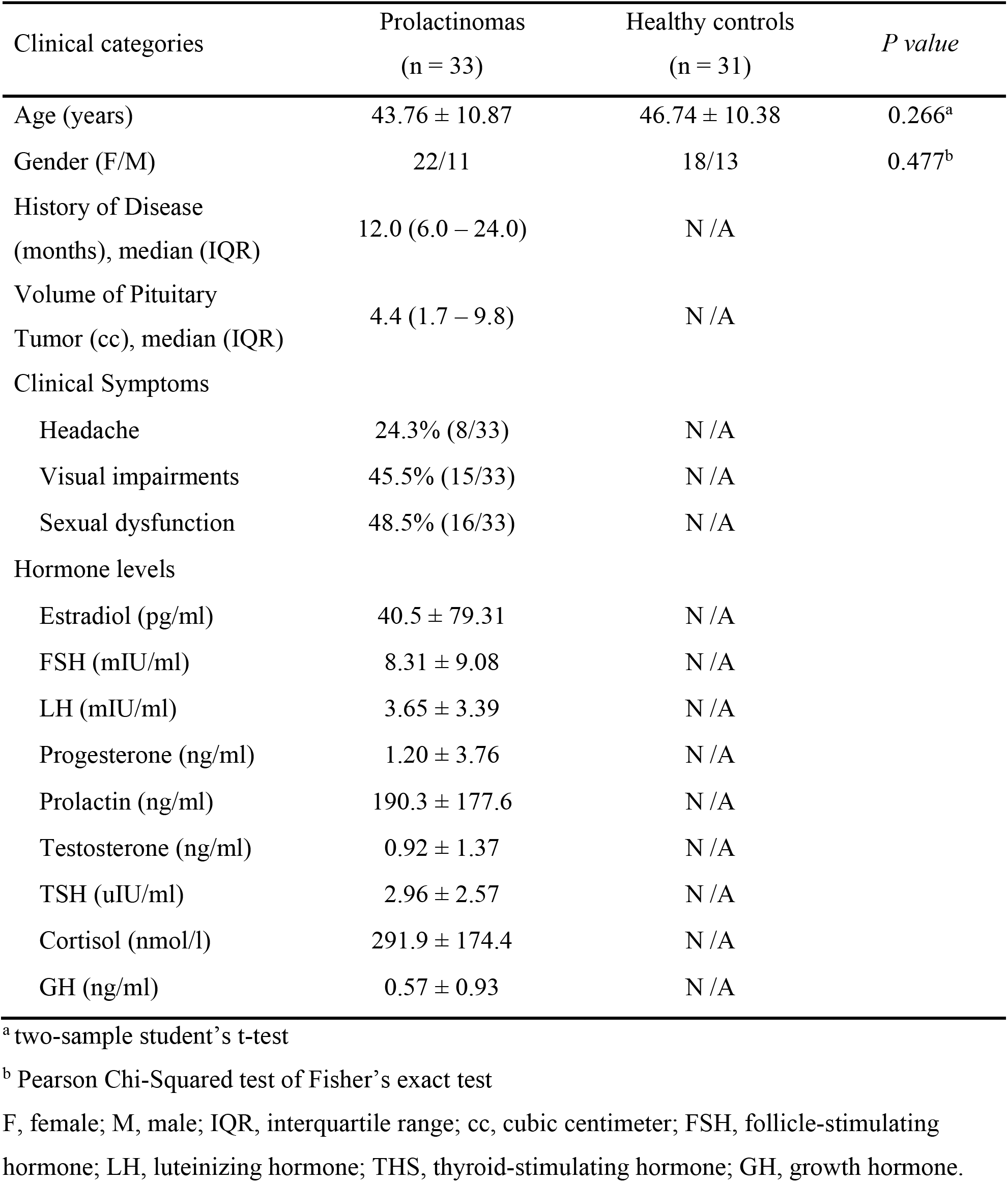
Clinical Features Comparison between Prolactinomas and Healthy Controls.

| Clinical categories | Prolactinomas<br>(n = 33) | Healthy controls<br>(n = 31) | *P value* |
| --- | --- | --- | --- |
| Age (years) | 43.76 ± 10.87 | 46.74 ± 10.38 | 0.266<sup>a</sup> |
| Gender (F/M) | 22/11 | 18/13 | 0.477<sup>b</sup> |
| History of Disease<br>(months), median (IQR) | 12.0 (6.0 – 24.0) | N /A |  |
| Volume of Pituitary<br>Tumor (cc), median (IQR) | 4.4 (1.7 – 9.8) | N /A |  |
| Clinical Symptoms |  |  |  |
| Headache | 24.3% (8/33) | N /A |  |
| Visual impairments | 45.5% (15/33) | N /A |  |
| Sexual dysfunction | 48.5% (16/33) | N /A |  |
| Hormone levels |  |  |  |
| Estradiol (pg/ml) | 40.5 ± 79.31 | N /A |  |
| FSH (mIU/ml) | 8.31 ± 9.08 | N /A |  |
| LH (mIU/ml) | 3.65 ± 3.39 | N /A |  |
| Progesterone (ng/ml) | 1.20 ± 3.76 | N /A |  |
| Prolactin (ng/ml) | 190.3 ± 177.6 | N /A |  |
| Testosterone (ng/ml) | 0.92 ± 1.37 | N /A |  |
| TSH (uIU/ml) | 2.96 ± 2.57 | N /A |  |
| Cortisol (nmol/l) | 291.9 ± 174.4 | N /A |  |
| GH (ng/ml) | 0.57 ± 0.93 | N /A |  |

<sup>a</sup> two-sample student's t-test
<sup>b</sup> Pearson Chi-Squared test of Fisher's exact test
F, female; M, male; IQR, interquartile range; cc, cubic centimeter; FSH, follicle-stimulating
hormone; LH, luteinizing hormone; TSH, thyroid-stimulating hormone; GH, growth hormone.

### Altered Whole-brain FC Patterns

Compared to healthy controls, prolactinomas showed increased FC between the left thalamus and visual cortex/association areas [bilateral lingual gyrus (LG), right intra-calcarine cortex (ICC), right supra-calcarine cortex (SCC), right cuneal cortex, and bilateral lateral occipital cortex (LOC)] and right cerebellum (Cereb), the right cerebellum and cerebral cortex [precuneus, posterior cingulate gyrus (PC), bilateral temporal fusiform cortex (TFusC)], and the left supplemental motor area (SMA) and right lingual gyrus (**Fig. 1**). In addition, the increased FC of the thalamus and visual cortex were more prominent in patients with visual field impairments (VFI) compared to patients without VFI (Supplementary Table 1).

**Figure 1.**
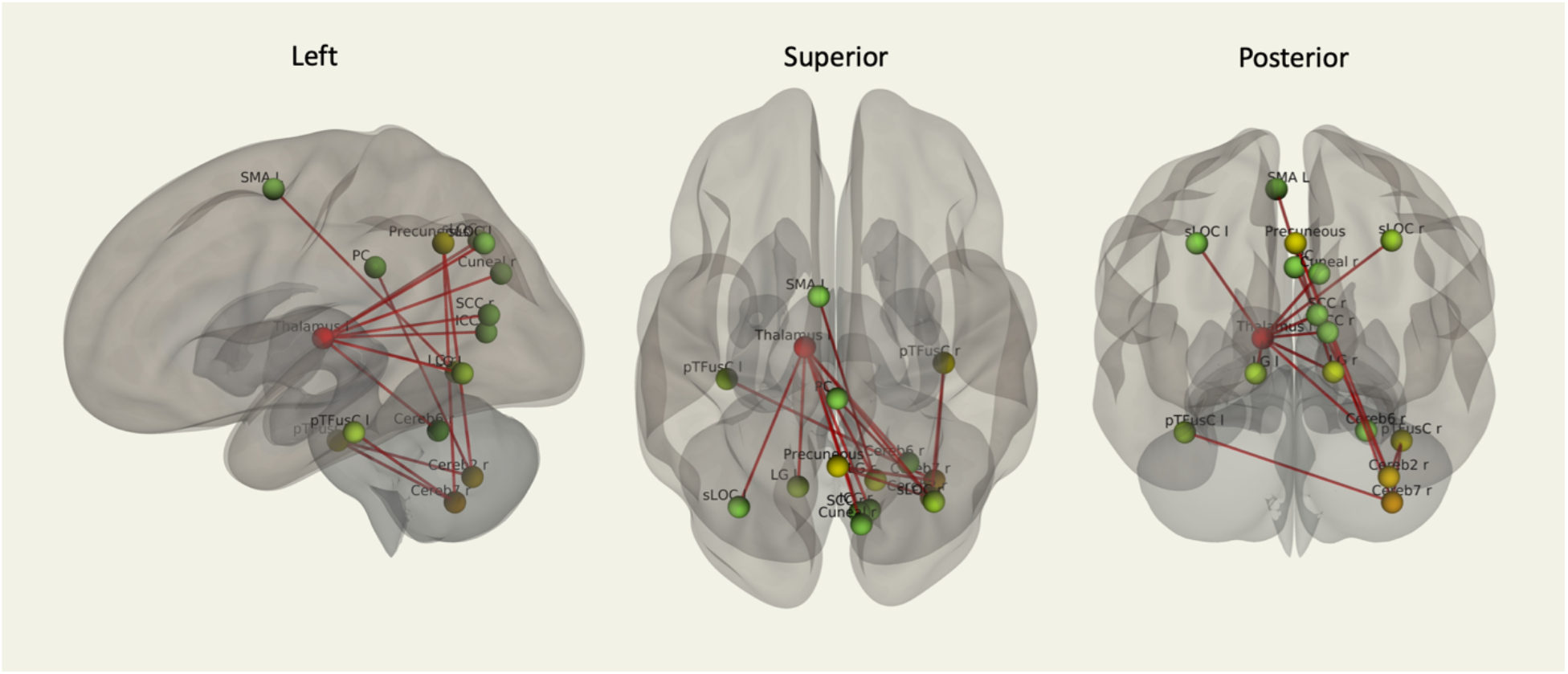
ROI-to-ROI analyses of the resting-state functional connectivity across the whole-brain. Compared to healthy controls, significantly increased functional connectivity was only shown in patients with prolactinomas. The false discovery rate (FDR) method was used to perform the seed-level two-sided correction at a significant level of *p* < 0.05. The nodes rendered by the red color indicated more connections to other nodes where the nodes rendered by the green color indicated less connections to other nodes, and the yellow-rendered nodes indicate mediate number connections in the whole connectivity pattern. Abbreviations: Cereb, Cerebellum; pTFusC, temporal fusiform cortex, posterior division; LG, lingual gyrus; sLOC, lateral occipital cortex, superior division; ICC, intra-calcarine cortex; SCC, supra-calcarine cortex; PC, cingulate gyrus, posterior division; SMA, supplemental motor area.

### Relationship between Hormone Levels and FC Metrics

In prolactinomas, females showed a significantly positive correlation between the prolactin level and the FC of left thalamus and right lingual gyrus. Males showed a positive correlation between the testosterone level and the FC of right cerebellum and left fusiform gyrus as well as the correlation between the LH level and the FC of left SMA and right lingual gyrus (**Fig. 2**).

**Figure 2.**
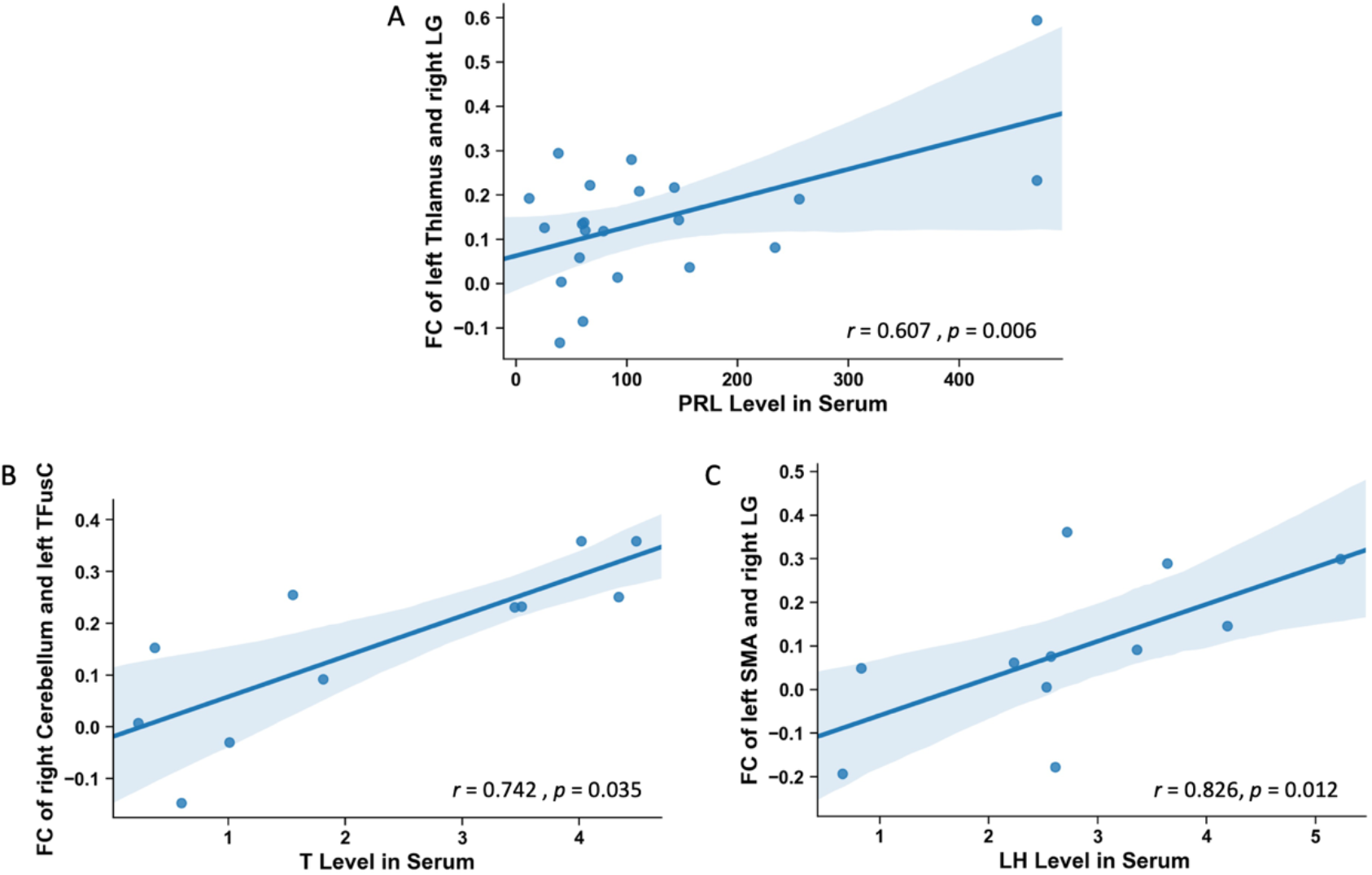
Partial correlations analyses between endogenous hormone levels and altered functional connectivity metrics. (A) the PRL level in serum is positively correlated with the FC of the left thalamus and right LG. (B) the T level in serum is positively correlated with the FC of the right cerebellum and left TFusC. (C) the LH level in serum is positively correlated with the FC of the left SMA and right LG. The shading area indicates the 95% confidence interval of a correlation line. Abbreviations: PRL, prolactin; T, testosterone; LH, luteinizing hormone; LG, lingual gyrus; TFusC, temporal fusiform cortex; SMA, supplemental motor area.

## DISCUSSION

To our knowledge, this study is the first to demonstrate the altered whole-brain FC patterns in prolactinomas and their relationships with endogenous hormone levels.

Our first notable finding is the illustration of increased functional connectivity between the left thalamus and visual cortex/association areas (lingual gyrus, ICC, SCC, cuneal cortex, and LOC). The thalamus is both the primary integrator and the gateway of sensory input to the cerebral cortex. Visual cortex/association areas mainly receive input of visual signals coming from the retina via the lateral geniculate nucleus of the thalamus. Considering the different visual-related FC patterns between patients with VFI and without VFI, these hyperactive intrinsic connections may illustrate compensatory activity within the visual system, which is consistent with a previous study.^3^ Another important finding is the increased cerebellar-cerebral connectivity that is mainly between the right cerebellum to the default mode network (DMN)^4^ (posterior cingulate cortex (PCC)/precuneus), and to bilateral TFusC. The PCC/precuneus plays a pivotal role in controlling the state of arousal and maintaining attention.^4^ TFusC mediates a variety of higher cognitive and emotional functions that relate to the processing of face/object perception.^5^ The cerebellum is also prominently activated in control of negative emotional processing and occurred concomitantly with the mirror neuron system, including TFusC and precuneus.^6^ Thus, there might exist functional connections between the cerebellum and DMN/TFusC. In addition, the thalamus plays a critical role in modulating the communication within cortico-cerebello-cortical parallel closed loops that involve DMN, salience, attention, and sensorimotor networks.^7^ Therefore, the increased FC of cerebellum and fusiform/DMN/thalamus may contribute to the hyperactive intrinsic connections serving dysfunctional cognitive or emotional processing in prolactinomas.

Hormones affect brain development, organization, and plasticity.^8^ The prolactin receptor is distributed broadly in the brain and can mediate the regulation of neuronal excitability, neurotransmission, and channels in the nervous system.^9^ Evidence has emerged supporting the critical role of tuberoinfundibular dopamine neurons in the remarkable plasticity and the necessity of their function during lactation.^10^ However, owing to the anti-correlation of prolactin and dopamine, the underlying mechanism of how prolactin influences brain reorganization is unknown. LH and testosterone may contribute to brain reorganization by modulating neural plasticity through routes engaging actin sytoskeleton^11^ and the growth of white matter in the brain.^12^ Thus, our findings of the relationships between hormone levels and the altered FC may illustrate the potential influence of endogenous hormones in brain reorganization in prolactinoma patients who usually display a cascade of hormone disorders.

However, several limitations should be addressed. First, the mass effect of the bulky tumor on surrounding structures was challenging to be completely ruled out even the tumor size was adjusted during the data analysis. Second, diffusion tensor imaging could be employed to assess the structural integrity of the visual pathway in future work. Third, the relationship between hormone levels and the altered FC in males might have been biased due to the relatively small sample size.

Overall, our findings provide new insights into the altered FC patterns and the important role of endogenous hormones in the neural mechanism of brain reorganization in prolactinomas.

## FUNDING

This work is supported by the funding from the National Science Foundation of China (NSFC) through grants: No.81571049 and No.81870863 and the National Institutes of Health (NIH) through Grants: R21CA198740, P41EB015898, P41RR019703, and R25CA089017. Shun Yao received the scholarship from the State Scholarship Fund, China Scholarship Council (201808440461).

## AUTHOR CONTRIBUTIONS

G.X. and J.S. provided the major research funding. S.Y., A.J.G., G.X., Y.T., and J.S. contributed to the conception and design of the study. S.Y. collected data, performed the data analysis, and drafted the manuscript. P.L. and C.C. did the brain tumor segmentation and J.S. reviewed all lesion masks (all of them were blind to the clinical data). P.L., P.J., W.V., A.Z., A.J.G., and R.Z participated in the fMRI data preprocessing and edited the manuscript. Critically revising the article: all authors. Approved the final version of the manuscript on behalf of all authors.

## DISCLOSURE

The authors declare no conflicts of interest.

## REFERENCES

1. Yao S, Song J, Gao J, et al. Cognitive Function and Serum Hormone Levels Are Associated with Gray Matter Volume Decline in Female Patients with Prolactinomas. Front Neurol [online serial]. 2018;8. Accessed at: https://www.frontiersin.org/articles/10.3389/fneur.2017.00742/full#. Accessed February 2, 2018.

2. van de Ven VG, Formisano E, Prvulovic D, Roeder CH, Linden DEJ. Functional connectivity as revealed by spatial independent component analysis of fMRI measurements during rest. Hum Brain Mapp. 2004;22:165–178.

3. Qian H, Wang X, Wang Z, Wang Z, Liu P. Altered Vision-Related Resting-State Activity in Pituitary Adenoma Patients with Visual Damage. PLoS ONE. 2016;11:e0160119.

4. Fransson P, Marrelec G. The precuneus/posterior cingulate cortex plays a pivotal role in the default mode network: Evidence from a partial correlation network analysis. NeuroImage. 2008;42:1178–1184.

5. Weiner KS, Zilles K. The anatomical and functional specialization of the fusiform gyrus. Neuropsychologia. 2016;83:48–62.

6. Schraa-Tam CKL, Rietdijk WJR, Verbeke WJMI, et al. fMRI Activities in the Emotional Cerebellum: A Preference for Negative Stimuli and Goal-Directed Behavior. Cerebellum. 2012;11:233–245.

7. Habas C, Manto M, Cabaraux P. The Cerebellar Thalamus. Cerebellum. 2019;18:635–648.

8. Garcia-Segura LM, Melcangi RC. Steroids and glial cell function. Glia. 2006;54:485–498.

9. Patil MJ, Henry MA, Akopian AN. Prolactin receptor in regulation of neuronal excitability and channels. Channels (Austin). 2014;8:193–202.

10. Barth C, Villringer A, Sacher J. Sex hormones affect neurotransmitters and shape the adult female brain during hormonal transition periods. Front Neurosci [online serial]. 2015;9. Accessed at: https://www.frontiersin.org/articles/10.3389/fnins.2015.00037/full. Accessed February 13, 2019.

11. Edwards BS, Clay CM, Ellsworth BS, Navratil AM. Functional Role of Gonadotrope Plasticity and Network Organization. Front Endocrinol [online serial]. 2017;8. Accessed at: https://www.frontiersin.org/articles/10.3389/fendo.2017.00223/full. Accessed July 13, 2019.

12. Perrin JS, Hervé P-Y, Leonard G, et al. Growth of White Matter in the Adolescent Brain: Role of Testosterone and Androgen Receptor. J Neurosci. 2008;28:9519–9524.

