## Supplemental Table 1. for "Hormone Levels Are Related to Altered Functional Connectivity in Prolactinomas"

**Supplementary Table 1.** Comparison of the altered functional connectivity related to the visual system in patients with and without VFI.

| **FC Metric** | **Non-VFI Patients** | **VFI Patients** | **Statistic** | ***p*-value** |
| --- | --- | --- | --- | --- |
| THA_L-sLOC_R | -0.044 ± 0.124 | -0.030 ± 0.148 | -0.308 | 0.760 |
| THA_L-sLOC_L | 0.070 ± 0.116 | 0.105 ± 0.127 | -0.815 | 0.422 |
| THA_L-LG_L | 0.181 ± 0.135 | 0.083 ± 0.147 | 1.988 | 0.056 |
| THA_L-LG_R | 0.157 ± 0.140 | 0.065 ± 0.135 | 1.889 | 0.068 |
| THA_L-ICC_R | 0.181 ± 0.137 | 0.053 ± 0.138 | 2.647 | 0.013* |
| THA_L-SCC_R | 0.125 ± 0.157 | 0.033 ± 0.117 | 1.857 | 0.073 |
| THA_L-Cuneal_R | 0.012 ± 0.199 | -0.037 ± 0.115 | 0.844 | 0.405 |

**Two-samples T-Test**

* *p* < 0.05

The two-sample student’s t-test was performed to do the comparison between groups.

Abbreviations: VFI, visual field impairment; THA, thalamus; LOC, lateral occipital cortex; LG, lingual gyrus; ICC, intra-calcarine cortex; SCC, supra-calcarine cortex; L, left; R, right.
